# Conditional forest models built using metagenomic data could accurately predict *Salmonella* contamination in Northeastern streams

**DOI:** 10.1101/2022.07.11.499664

**Authors:** Taejung Chung, Runan Yan, Daniel L. Weller, Jasna Kovac

## Abstract

The use of water contaminated with *Salmonella* for produce production contributes to foodborne disease burden. To reduce human health risks, there is a need for novel, targeted approaches for assessing the pathogen status of agricultural water. We investigated the utility of water microbiome data for predicting *Salmonella* contamination of streams used to source water for produce production. Grab samples were collected from 60 New York streams in 2018 and tested for *Salmonella*. Separately, DNA was extracted from the samples and used for Illumina shotgun metagenomic sequencing. Reads were trimmed and used to assign taxonomy with Kraken2. Conditional forest (CF), regularized random forest (RRF), and support vector machine (SVM) models were implemented to predict *Salmonella* contamination. Model performance was determined using 10-fold cross-validation repeated 10 times to quantify area under the curve (AUC) and Kappa score. Taxa identified as the most informative for accurately predicting *Salmonella* contamination based on conditional variable importance were compared to taxa identified by ALDEx2 as being differentially abundant between *Salmonella*-positive and - negative samples. CF models outperformed the other two algorithms based on AUC (0.82 - CF, 0.76 - RRF, 0.67 - SVM) and Kappa score (0.41- CF, 0.38 - RRF, 0.19 - SVM). CF and differential abundance tests both identified *Aeromonas* (VI = 0.32) and *Tabrizicola* (VI = 0.12) as the two most informative taxa for predicting *Salmonella* contamination. The taxa identified in this study warrant further investigation as indicators of *Salmonella* contamination in Northeastern freshwater streams.

**IMPORTANCE:** Understanding the associations between surface water microbiome composition and the presence of foodborne pathogens, such as *Salmonella*, can facilitate the identification of novel indicators of *Salmonella* contamination. This study assessed the utility of microbiome data and three machine learning algorithms for predicting *Salmonella* contamination of Northeastern streams. The research reported here both expanded the knowledge on the microbiome composition of surface waters and identified putative novel indicators (i.e., *Aeromonas* and *Tabrizicola*) for *Salmonella* in Northeastern streams. These putative indicators warrant further research to assess whether they are consistent indicators of *Salmonella* for regions, waterways, and years not represented in the dataset used in this study.

## INTRODUCTION

According to US Centers for Disease Control and Prevention (CDC), 46% of foodborne illnesses in the US caused by a known food vehicle between 1998 and 2008 were linked to produce commodities consumption (1). In the US, *Salmonella* is the most common bacterial pathogen associated with outbreaks linked to fresh produce (2, 3). Thus, preventing *Salmonella* contamination of fresh produce is critical for managing foodborne disease burden in the US.

Multiple produce-associated outbreaks have been putatively traced back to the use of contaminated water for produce production (4–7). Therefore, identifying when water is likely to be contaminated is a central component of produce safety risk management plans. In many countries, agricultural and recreational water *E. coli*-based standards have been established (8–11). However, *E. coli* is an indicator of fecal and not pathogen contamination. Indeed, the presence and direction of the association between *E. coli* levels and foodborne pathogen presence varies substantially within the scientific literature, with some studies reporting positive relationships (12–16), and others reporting negative or no relationship (17–20). As a result, *E. coli* appears to be an unreliable indicator of *Salmonella* contamination of surface waterways, even though *E. coli* can be used as a general indicator of hygienic condition of water (17, 21). Thus, there is a need for novel approaches for identifying when and where agricultural waterways may be contaminated with foodborne pathogens, such as *Salmonella*.

Metagenomics opened new avenues for characterization of water microbiomes. Concurrent characterization of microbiome and pathogen status in water provides an opportunity for the identification of microbial taxa associated with pathogen contamination of agricultural water. Such taxa could be identified by developing models that use microbiome data (i.e., presence or absence of taxa, or differences in their relative abundance) to predict when and where pathogens are present. However, since the existing water microbiome literature demonstrates substantial spatial and temporal variation in water microbiome composition (22–25), identification of such “indicator” taxa is difficult using conventional analytical approaches, such as multivariate ordination (25). Machine learning provides an alternative approach that may be useful for identifying “indicator”, or combinations of “indicator” taxa and for developing classification models that use these taxa to predict pathogen contamination status (26, 27).

Among classification machine learning models, supervised models are particularly useful when the outcome information, such as pathogen status, is known for a set of samples. Labelling the training dataset with an outcome class label allows for the development of a classifier that can predict pathogen status. A variety of supervised classification models have been developed to address different data structure challenges and improve the accuracy of prediction (28–31). A benchmarking study that applied multiple machine learning models on human gut microbiome found differences in the performance of models based on different machine learning algorithms. This is likely due to differences in the characteristics of algorithms, such as linear/non-linear separation, ensemble or regression approaches (30). This suggests the importance of selecting and testing multiple algorithms to improve the prediction accuracy. In addition to model selection, the performance of a model may be affected by microbiome data pre-processing, such as data normalization (28). The latter is commonly carried out to account for potential differences in sample library sizes, hence its effect on prediction accuracy needs to be assessed (31–33).

With the above-outlined consideration in mind, we applied multiple machine learning classifiers to normalized and non-normalized microbiome data for samples collected from 60 different streams. Our goal was to identify microbial indicators predictive of *Salmonella* contamination in stream water samples collected in a region in Northeastern United States. Lastly, we also assessed whether the addition of data on physicochemical properties of water samples increases the accuracy of predicting *Salmonella* contamination.

## RESULTS

### Samples were sequenced with a median of 5,956,185 reads and a median of 8.955% reads were assigned bacterial taxonomic identifier

A median of 5,956,185 reads per sample were obtained from 60 samples [min = 4,048,684, max = 9,301,059, standard deviation (SD) = 1,125,759] and median of 8.95% reads were classified as bacterial using metagenomics taxonomic classifier Kraken2 [median = 529,963, min = 145,211, max = 1,059,311, standard deviation (SD) = 206,336]. Across samples, a total of 885 different genera from 307 different families were assigned.

### The overall microbiome composition was not associated with the presence of *Salmonella* in surface water samples

The principal component analysis (PCA) biplot (Fig. 1B and 1D) and scree plot (Fig. 1A and 1C) showed that the sample microbiomes did not cluster based on the presence of *Salmonella*. As evident from the scree plot (Fig. 1A and 1C), the first two components explained a relatively low percent of variance in the microbiome composition. Specifically, they explained 28.3% of the variance at the genus level (Fig. 1A) and 26.5% of the variance at the family level (Fig. 1C). PERMANOVA results also did not indicate significant association between microbiome composition and *Salmonella* isolation (p = 0.318 (family-level), p = 0.349 (genus-level)).

**FIG 1.**
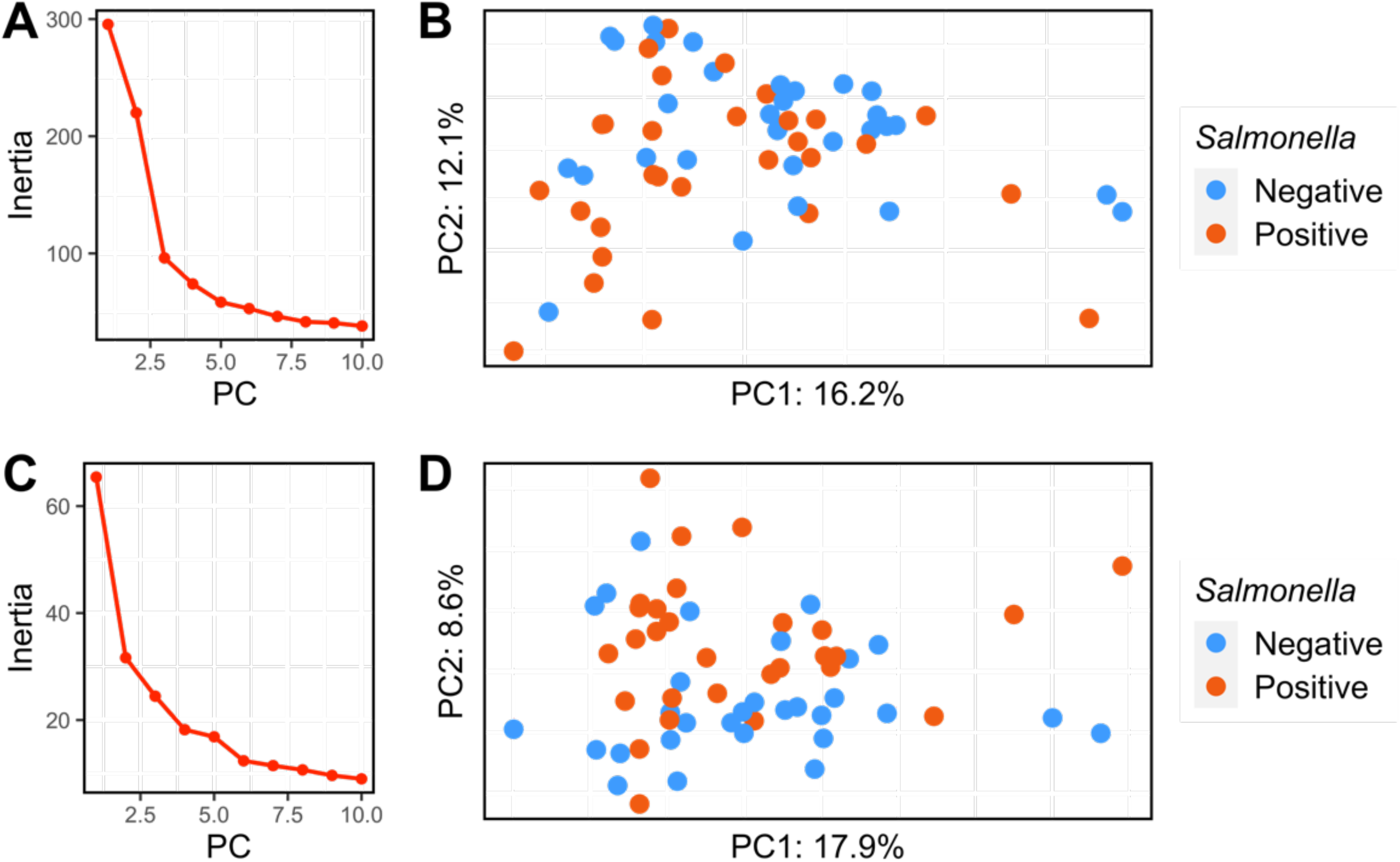
Principal Component Analysis (PCA) based on the Aitchison distance. Scree plot between principal components and eigen values are shown at the **(A)** genus level and **(C)** family level. The PCA biplot showing ordination of samples between based on the microbiome composition at the **(B)** genus level and **(D)** family level and color-coded based on whether *Salmonella* was detected (orange) or not detected (blue) in water samples.

### CF models outperformed RRF and SVM models at predicting *Salmonella* contamination

Regardless of feature set, the area under the curve (AUC) and Kappa score were always higher for conditional forest (CF) compared to regularized random forest (RRF) and support vector machine (SVM) when genus-level microbiome data were used (Fig. 2). Furthermore, the AUC was also consistently higher for CF models compared to RRF and SVM models when family-level microbiome data were used. However, the Kappa values were similar for CF and RRF models when family-level microbiome data were used (Fig. 2). RRF models (AUC = [0.68, 0.75], Kappa = [0.24, 0.35]) and SVM models (AUC = [0.61, 0.64], Kappa = [0.1, 0.15]) had lower AUC and Kappa score than CF models (AUC = [0.76, 0.82], Kappa = [0.32, 0.42]). We found that the AUC range for CF did not overlap with AUC ranges of other two methods, indicating that CF has outperformed RRF and SVM. Hence, further analyses were carried out using the CF models.

**FIG 2.**
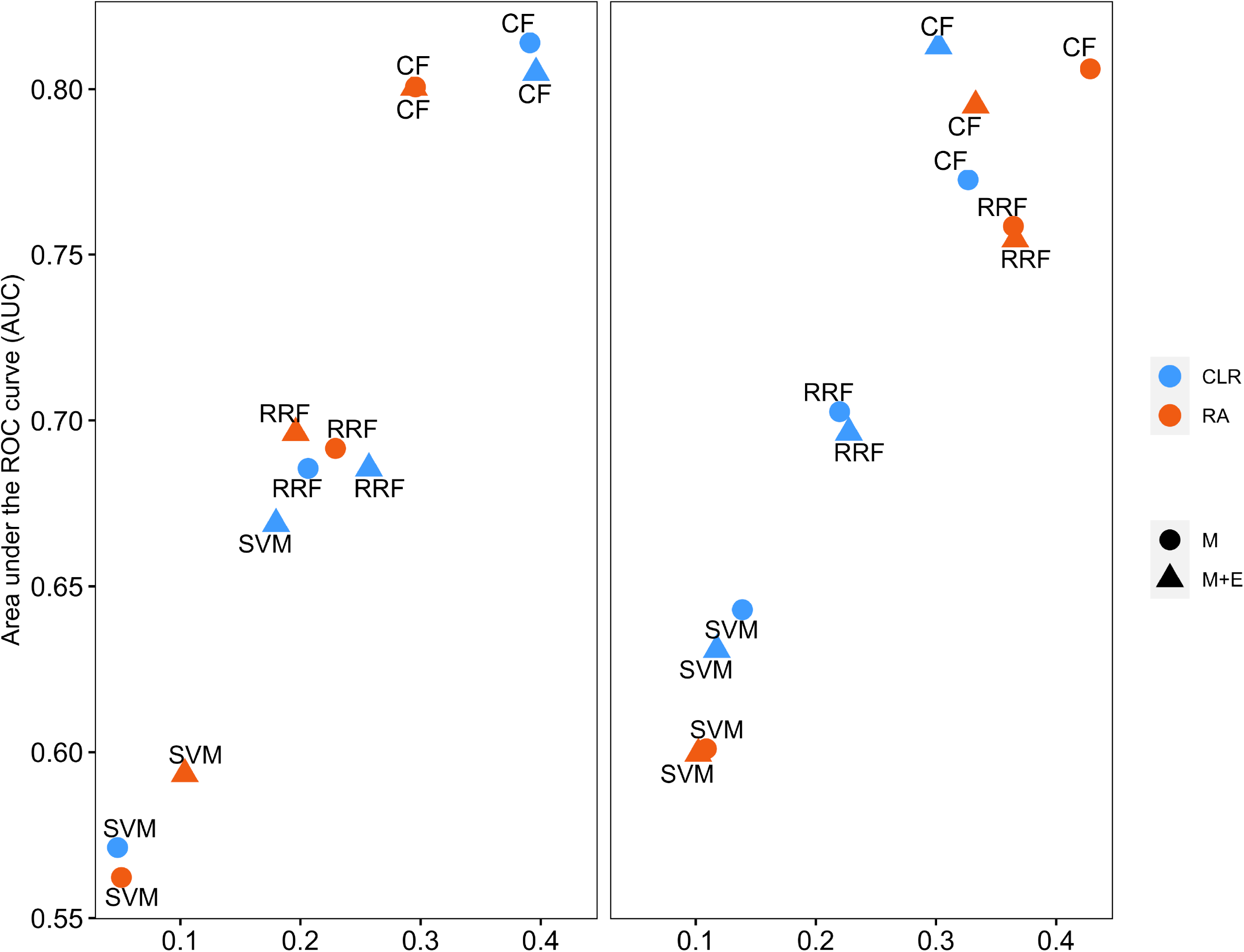
Kappa score and Area Under the Curve (AUC) for each machine learning algorithm. Results are shown at two taxonomic levels used for the classification (left – genus, right– family). Two different data transformation method (CLR [blue] – centered log-ratio transformation, and RA [orange] – relative abundance) were compared with two different data structures: M (circles) – microbiome data only and M+E (triangles) – microbiome and environmental data.

Across all models using the genus-level data, the CF model run on CLR-transformed relative abundance data without environmental features had the highest AUC (0.81) and Kappa score (0.42) (Fig. 2). When using family level data, the CF model using the relative abundance data, without environmental features resulted in the highest Kappa score (0.42), and second highest AUC (0.80) (Fig. 3).

**FIG 3.**
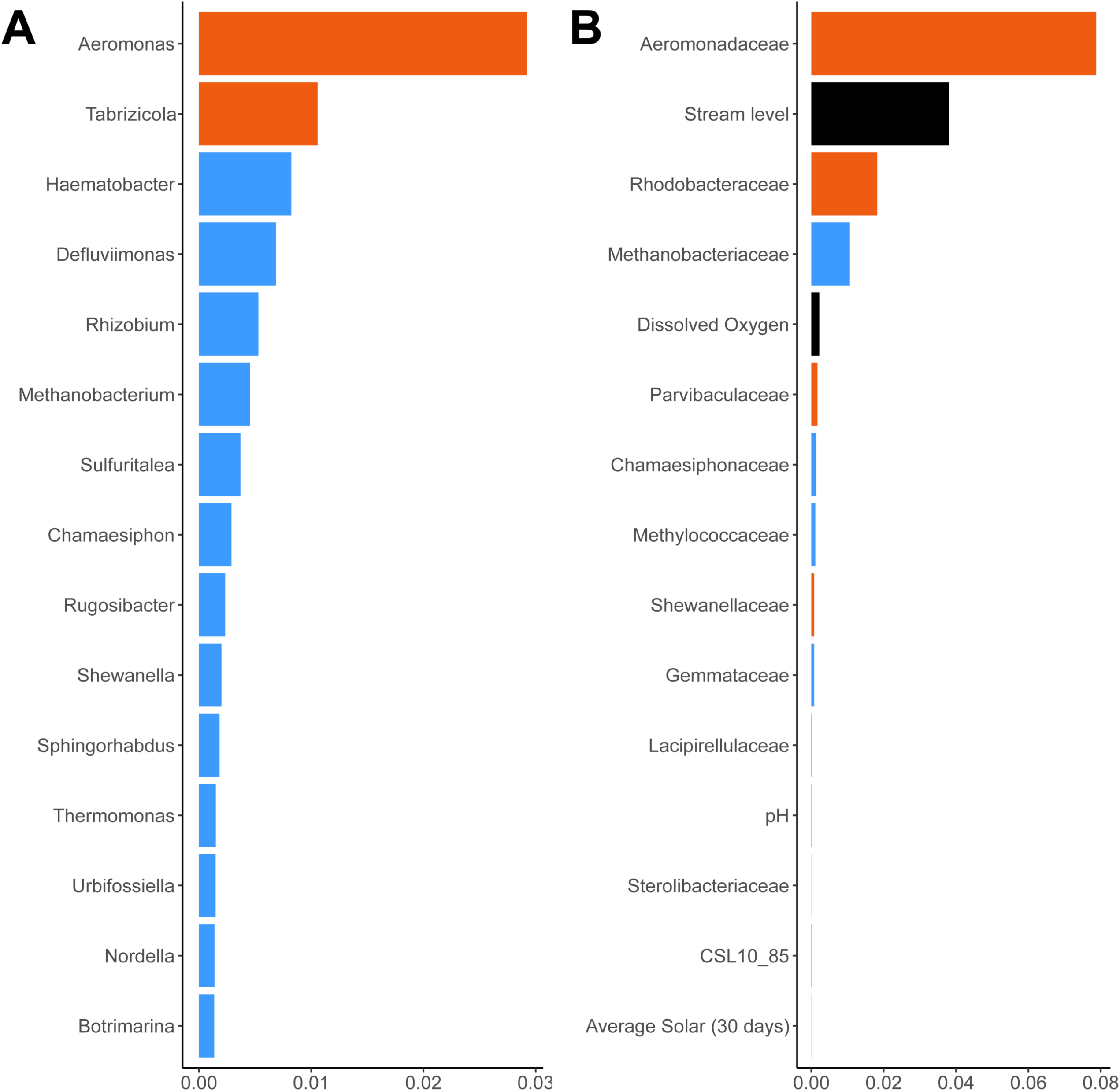
Conditional variable importance of taxa. Conditional variable importance was calculated from best performing conditional forest (CF) models of **(A)** genus level and **(B)** family level. Top 15 most informative for prediction of *Salmonella* contamination are presented. Orange bars indicate taxa significantly differentially abundant, blue bars indicate taxa not significantly differentially abundant, and black bars indicate environmental features (CSL 10_85 = change in elevation divided by the length between points 10 and 85 percent of distance along main channel to basin divide).

### *Aeromonas* and *Tabrizicola* were most informative for accurately predicting *Salmonella* presence

Using genus-level microbiome data, CF identified *Aeromonas, Tabrizicola, Haematobacter, Defluviimonas*, and *Rhizobium* as the five most informative genera for predicting *Salmonella* contamination (Fig. 3A). In family level analysis, *Aeromonadaceae, Rhodobacteraceae*, and *Methanobacteriaceae* were identified as the three most informative families. Furthermore, four environmental features (i.e., stream level, dissolved oxygen level, pH, and changes of elevation and length of the stream) were also identified as informative features in the CF model that included environmental features (Fig. 3B).

The differential abundance analysis carried out using ALDEx2 and Kruskal-Wallis test identified two bacterial genera, *Aeromonas* and *Tabrizicola*, (Fig. 4) as significantly differentially abundant between *Salmonella*-positive and *Salmonella-*negative samples. These two genera belong to *Aeromonadaceae* and *Rhodobacteraceae* families, respectively. These two families are among the families identified as differentially abundant between *Salmonella*-positive and -negative samples (i.e., *Aeromonadaceae, Parvibaculaceae, Rhodobacteraceae*, and *Shewanellaceae*) (Fig. 5).

**FIG 4.**
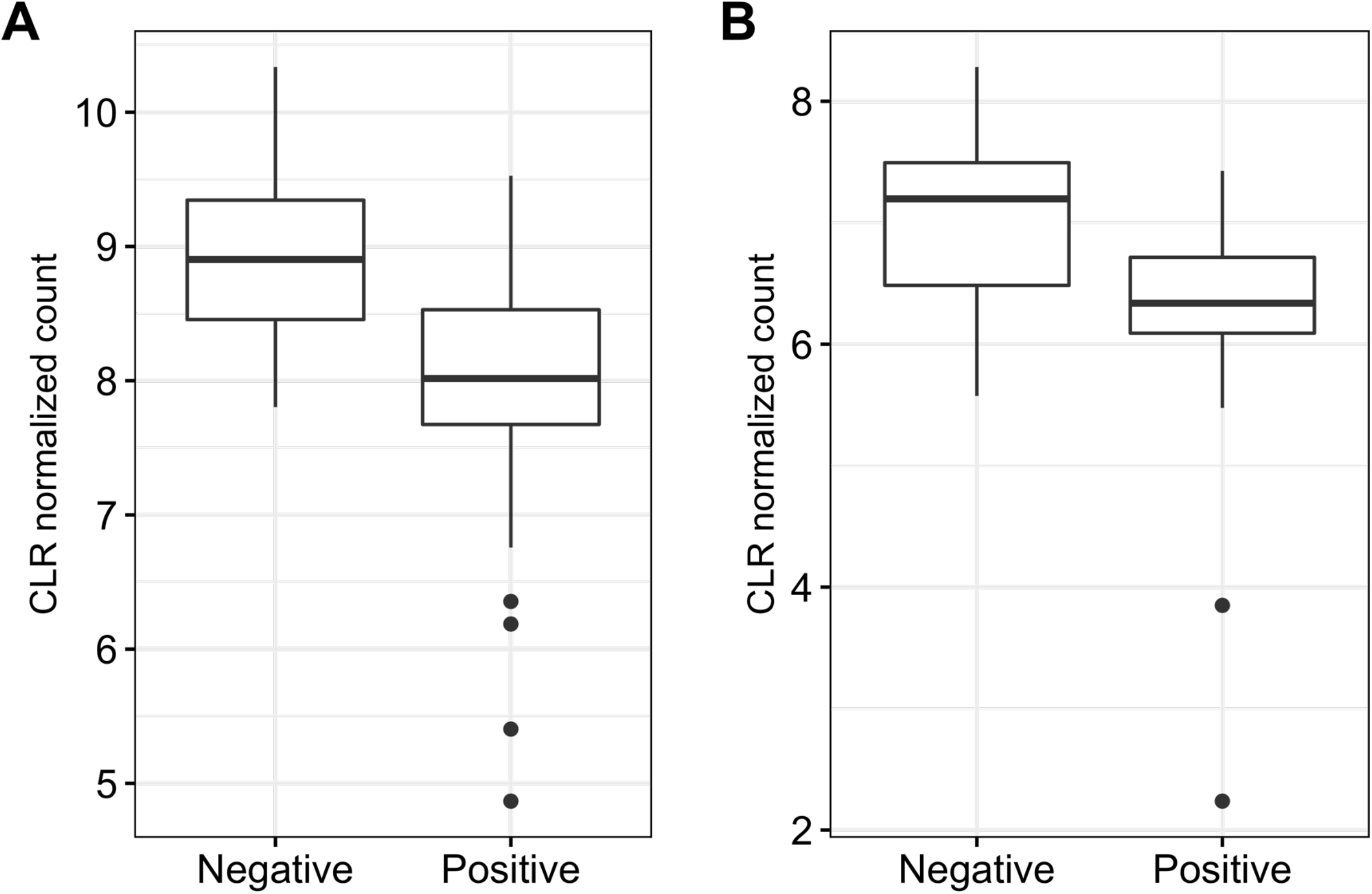
Relative abundance of **(A)** *Aeromonas* and **(B)** *Tabrizicola* in water samples. These genera were identified as significantly differentially abundant between *Salmonella* positive and negative water samples.

**FIG 5.**
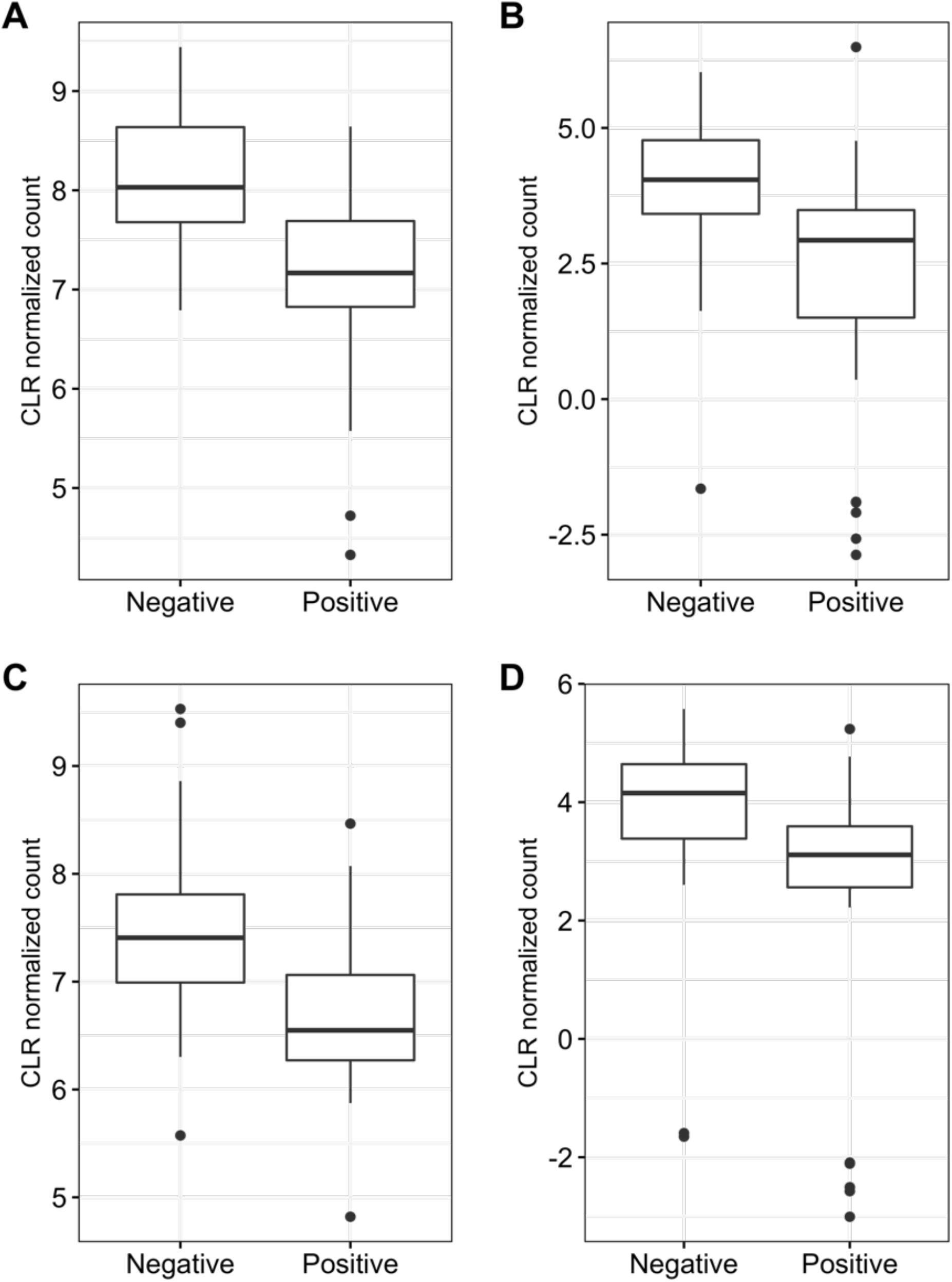
Relative abundance of **(A)** *Aeromonadaceae* **(B)** *Parvibaculaceae* **(C)** *Rhodobacteraceae*, and **(D)** *Shewanellaceae*. These families were identified as significantly differentially abundant between *Salmonella* positive and negative water samples.

## DISCUSSION

This study was based on the premise that contamination of surface waters with enteric pathogens such as *Salmonella*, co-occurs with other microbiota that is either linked with the contamination source or linked with the environmental conditions that favor pathogen introduction to or survival in freshwater systems. As such, these taxa could be used to identify when and where surface waters may by contaminated by *Salmonella*. Hence, we aimed to leverage water microbiome data and machine learning classifiers to identify specific taxa predictive of *Salmonella* contamination that could be further developed into rapid detection assays (10). For example, taxa that (i) consistently occur in samples contaminated with *Salmonella* or are (ii) consistently present in a significantly different relative abundance in samples contaminated with *Salmonella*, may be utilized to develop rapid PCR-based indicator assays.

### Data transformation did not have a notable effect on the performance of predictive models

Microbiome data analyses are challenging due to inherent data complexities. These include data sparsity (i.e., many taxa are only present in small proportions of samples, resulting in a large proportion of zero counts), collinearity (i.e., some taxa are highly correlated), imbalanced library sizes, and a well-known “small n large p” problem (i.e., small number of samples and a large number of taxa). Some of these challenges can be addressed prior to applying machine learning models or can be addressed by applying machine learning methods that can address certain microbiome data challenges. Here, we used a central log-ratio (CLR) transformation to mitigate the potential effects of different library sizes, and to take into consideration the compositional nature of microbiome data (33, 34). We found that the microbiome data transformation had an inconsistent effect on a model performance, as it improved the AUC of some models (e.g., CF using genus- and family-level data, SVM using family-level data) while it decreased the AUC of other models (e.g., RRF using genus- and family-level data, SVM using genus-level data). Previous studies on gut microbiome data emphasized the importance of data transformation due to the compositionality of the microbiome data (28, 30, 32). However, they also reported that the performance of tree-based algorithms (i.e., random forest and XGBoost) was not significantly affected by data transformation (32).

### Model selection is critically important when applying machine learning to microbiome data

Different machine learning classifiers address microbiome data challenges to various degrees; hence we assessed the performance of three different machine learning algorithms for prediction of *Salmonella* contamination. At both genus-level and family-level classification, we found that CF performed overall better than RRF and SVM. This is consistent with other studies that compared multiple algorithms for predicting foodborne pathogen contamination of surface water, and consistently found that CF was a high performing algorithm (27). However, many of these studies also found that RRF and/or SVM also performed well. Given the highly correlated nature of microbiome data, it is not surprising that CF, which was developed to address limitations of other random forest algorithms for handling correlated data, outperformed the RRF and SVM (35–37). Moreover, given the complex relationships that underpin microbial ecosystems, hierarchical relationships between the presence-absence of various taxa (e.g., models that incorporate “interactions” or hierarchy such as CF and RRF) may predict pathogen presence better than algorithms that do not (e.g., SVM).

### Differential abundance analysis and conditional forest models identified putative indicators of *Salmonella* contamination in surface waters

When assessing the association between the overall microbiome composition and *Salmonella* contamination, associations between certain taxa and *Salmonella* contamination may be missed due to the large number of taxa included in the analyses. Indeed, we found the lack of association between the overall microbiome composition and the presence of *Salmonella* based on the Principal Component Analysis (PCA) and PERMANOVA. Hence, we used differential abundance analysis and machine learning algorithm CF to discover individual taxa (or their relative abundance) associated with *Salmonella* contamination.

Both, differential abundance analysis and CF identified some of the same taxa predictive of *Salmonella* contamination in stream water samples. Using ALDEx2 differential abundance analysis and CF, we identified two bacterial genera (*Aeromonas* and *Tabrizicola*) which were present in a significantly lower relative abundance in samples contaminated with *Salmonella. Aeromonas* species have previously been found in natural water and a broad range of foods, in addition to human and animal gastrointestinal tracts (38, 39). *Aeromonas* is regarded not only as an important pathogenic bacterium in fish and cold-blooded animals, but also as an opportunistic pathogen in immunocompromised humans (38). *Aeromonas* (belonging to *Aeromonadaceae*) has similar morphological and biochemical characteristics as *Enterobacteriaceae*, a family of microorganisms commonly used as an indicator of poor hygienic conditions in food systems (40). Bonadonna et el. reported that the presence of *E. coli* and fecal coliforms were associated with lower counts of *Aeromonas*, whereas the prevalence of total coliforms was associated with higher counts of *Aeromonas* in bathing waters along the sea-coast of the Adriatic Sea (41, 42). Another identified genus, *Tabrizicola*, belongs to a family of *Rhodobacteraceae*, which is usually found in aquatic environment, including lakes and wastewater treatment facilities (43–45). However, the ecological role of *Tabrizicola* is still understudied. In addition to the two genera discussed above, we identified several bacterial families positively and negatively associated with *Salmonella* contamination in water. Four families were identified in both differential abundance analysis and machine learning variable importance analysis (i.e., *Aeromonadaceae, Rhodobacteraceae, Shewanellaceae*, and *Parvibaculaceae*). These four families are known marine or aquatic microbiome members commonly found in natural waters (46–48). However, their relationship with *Salmonella* is unknown. A study by Gu et al. used 16S rDNA amplicon sequencing data and found an association between specific microbial taxa and the prevalence and population density of *Salmonella enterica* detected in ponds and wells in Eastern Shore of Virginia (ESV) between January to December (49). They found that the relative abundance of *Sphingomonadales* was significantly correlated with *S. enterica* prevalence as well as its population density in irrigation ponds and water wells. However, in our study, *Sphingomonadales* were not identified as informative for prediction of *Salmonella* contamination using machine learning nor differential abundance analysis (49). This inconsistency could potentially be explained by regional differences in the water microbiome composition, which is known to be influenced by environmental features such as natural variation over time, weather, and adjacent land use (22, 23, 25).

In this study, we found that several environmental features were informative for classification in the CF model. CVI of stream level was the second most informative feature for predicting *Salmonella* contamination, and level of dissolved water, pH, and change in elevation and length (CSL 10_85) were weakly associated with *Salmonella* contamination. Previous study reported that environmental features had an effect on the level of *E. coli* and the probability of detecting foodborne pathogen from the fresh water samples (20). Therefore, it is important to consider environmental condition as supplementary features when developing new tools for predicting *Salmonella* contamination of fresh water based on the specific taxa.

Some of the differences between taxa identified in this study and previous studies may be due to the differences in microbiome composition between surface, pond, and well waters. Thus, future studies that sample the same (or multiple) water types over multiple growing regions from multiple states are needed to assess whether the putative indicators of *Salmonella* contamination identified in this study are reproducible more broadly and are suitable candidates for the development of a rapid nucleic acid-based diagnostic assay.

## Conclusions

This study applied machine learning classifiers and differential abundance analyses of surface water microbiome data to identify putative novel indicators of *Salmonella* contamination. We identified *Aeromonas* and *Tabrizicola* bacterial genera and *Aeromonadaceae, Rhodobacteraceae, Shewanellaceae*, and *Parvibaculaceae* families that warrant further assessment as putative indicators of *Salmonella* contamination of the water. The identified taxa are potential targets for the development of an alternative or complementary (to *E. coli* quantification) water quality/safety monitoring strategy focused on mitigating the use of surface waters likely contaminated with *Salmonella*. However, the models developed in this study first need to be validated on new samples collected from a broader geographic area and over multiple seasons to assess the predictive accuracy of taxa identified here. Furthermore, deeper metagenomic sequencing that would enable metagenome assembly may facilitate identification of putative novel indicators of *Salmonella* contamination at a species level, as well as characterization of their functional potential.

## MATERIALS AND METHODS

### Sample collection and processing

Water samples were collected from sixty streams in Upstate New York state between July and October 2018 as described by Weller et al. (2020) (20). All chemical, microbial, and environmental water quality data were previously reported by Weller et al. (2020) (20). Briefly, 10 L grab samples were collected from each stream and tested for *Salmonella* presence. Each grab sample was filtered through modified Moore swabs (mMS). Buffered peptone water supplemented with novobiocin (20 mg/l) was added to Whirl-Pak bags containing mMS and incubated at 35°C for 24 h. After incubation, a BAX real-time PCR screen was used to identify samples that were presumptively positive for *Salmonella. Salmonella* presence was confirmed using culture-based methods fully described in Weller et al. (20)

Separately, 100 ml grab samples were collected for metagenomic analysis. The 100-mL samples were filtered through a 0.45 mm filter (Nalgene, Thermo Fisher Scientific, Waltham, MA USA). Filters were then stored at -80 °C until DNA extraction.

### DNA extraction and microbiome sequencing

DNA was extracted using DNeasy Power Water kit (Qiagen, MD, USA) per manufacturer’s instructions. Extracted DNA was examined for quality and quantified using Nanodrop One (Thermo Fisher Scientific, MA, USA) and Qubit 3 (Thermo Fisher Scientific, MA, USA), respectively. DNA was then sent to the Penn State Genomics Core Facility for library preparation and sequencing. Libraries were prepared using Nextera XT Flex per manufacturer’s instructions. Pooled libraries were sequenced on an Illumina NextSeq with 150 bp paired end reads.

### Sequence quality control and taxonomic classification

FastQC version 0.11.5 was used to assess read quality using default parameters (50). Illumina adapters and low-quality bases were trimmed using Trimmomatic (v 0.36) (51) with default parameters. Trimmed reads were taxonomically classified using Kraken2 (v 2.1.2) (52) and relative abundances inferred using Bracken (v 2.5) (53). The NCBI’s RefSeq nucleotide database (v 207) (54) was used to build a Kraken2 database. Any read that mapped to a single reference genome was labeled with the NCBI taxonomic annotation (taxid) corresponding to that reference genome. Any read that mapped to multiple reference genomes, or did not meet or exceeded the confidence scoring threshold was assigned a last common ancestor (LCA) taxonomic identification (taxid) (52).Confidence scores were set to 0.1, meaning that at least 10% of the total number of kmers from a read were classified. Bracken was used to estimate the abundance of taxa by re-distributing reads in the taxonomy using Bayes’ theorem (53). Assigned taxonomy and taxid counts of all samples were merged into a table that was used for downstream analyses.

### Microbiome Analyses

All statistical analyses of microbiome data were performed in R (version 4.1.0; R core Team, Vienna, Austria) (55), using a compositional analyses framework (33). First, the estimated abundances were transformed using the centered log-ratio (CLR) transformation (34). Ratio transformations capture the relationship between the taxonomic units in the data, and logarithm of these ratios ensures that the data are symmetric and linearly related (34). Distances between samples were calculated using the Aitchison distance (i.e., Euclidian distance after CLR transformation) to investigate the among-sample differences in microbiome composition [11]. Principle component analysis (PCA) was carried out using the ‘princomp’ function in R on relative abundances of taxids to visualize the ordination and clustering of samples based on the microbiome composition (33). The first two principal components were plotted using the ‘ggplot2’ package (v 3.3.3) (56). Samples were color-coded to visually assess whether they cluster based on the *Salmonella* presence/absence. Permutational Multivariate Analysis of Variance (PERMANOVA) was carried out to assess statistical associations between microbiome composition and *Salmonella* presence using the ‘adonis’ function in the ‘vegan’ package (v 2.5.7) (57). Differential abundance test was conducted using the ALDEx2 R package (58) to identify bacterial genera and families that were differentially abundant between *Salmonella*-positive and - negative samples. Each identified bacterial genus and family were tested using Kruskal-Wallis test to assess statistical significance of detected differences in their relative abundance (59).

### Predictive modelling

Three machine learning algorithms (i.e., conditional forest (CF) (35), regularized random forest (RRF) (36), and support vector machine (SVM) with sigmoid kernel (37)) that had previously been applied on microbiome data (60–62) and were suitable for microbiome data structure (27) were used in this study. Additionally, these algorithms were previously reported to outperform others for predicting *Salmonella* presence using environmental data (27). These methods were used to develop models that predict *Salmonella* presence or absence in water samples. Both relative abundance of taxid and CLR transformed relative abundance of taxid were separately used as features to assess the effect of microbiome data transformation on model performance. Model training and evaluation was performed using the ‘mlr’ package (v 2.19.0) (63). Ten-fold cross-validation repeated three-times was used to tune hyperparameters to maximize area under the curve (AUC) (64, 65). In total, analyses were conducted using two feature sets: (i) untransformed relative abundances of microbial taxa and (ii) CLR-transformed relative abundances of microbial taxa, to assess whether models perform better on transformed microbiome data. Analyses were also carried out at two different taxonomic levels (i.e., genus and family) and with or without environmental data. In total, 24 models were constructed separately based on the AUC and kappa scores (Table 1). The best performing model for each combination of algorithm and input data (Table 1) was selected for identifying informative taxa. conditional variable importance (CVI) was calculated using the ‘party’ package (v. 1.3.7) for CF models (35).

**TABLE 1.**
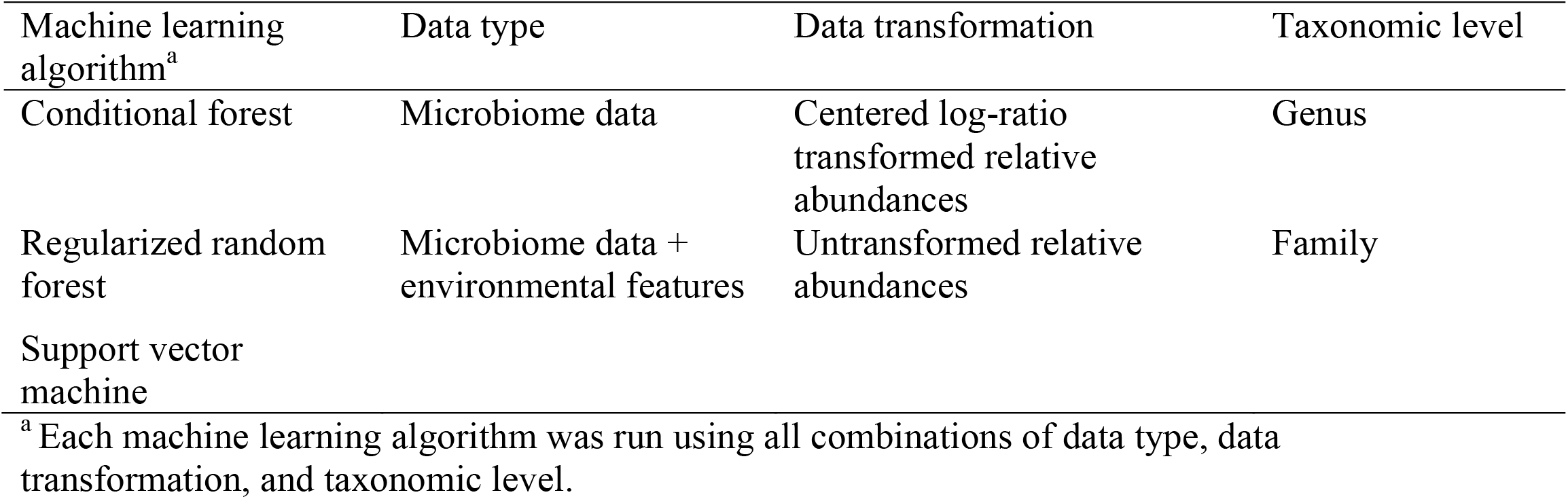
Machine learning algorithms, data types, data transformation, and taxonomic levels used for prediction of *Salmonella* contamination.

## Acknowledgments

This work was supported by the Penn State Institutes of Energy and the Environment seed grant, the Penn State Huck Institutes of the Life Sciences’ Genomics Core facility, and the USDA National Institute of Food and Agriculture Hatch Appropriations under Project #PEN04646 and Accession #1015787. We thank Martin Wiedmann for the support of DW’s work at the Cornell University.

## Data availability

Sequences generated in this study are available in the NCBI Sequence Read Archive database under the BioProject accession number PRJNA849616. Script used for bioinformatics and statistical analyses are available in GitHub repository: https://github.com/tuc289/SurfaceWaterMicrobiome/tree/master/Year2.

